# Antinociceptive synergy in a peripheral hyperalgesia model: interplay of cannabinoidergic, adrenergic, and opioidergic systems and their antagonism

**DOI:** 10.1101/2024.05.30.596637

**Authors:** Caio Fabio Baeta Lopes, Thamyris Santos-Silva, Flávia Cristina Fonseca, Bárbara Formiga Gonçalves de Queiroz, Igor Dimitri Gama Duarte, Thiago Roberto Lima Romero

**Affiliations:** Laboratory of Pain and Analgesia, Department of Pharmacology, Institute of Biological Sciences, Federal University of Minas Gerais, Belo Horizonte, Brazil

**Keywords:** isobolographic analysis, mechanical nociception, algesimetric test, prostaglandin-induced hyperalgesia, peripheral pain

## Abstract

There is growing interest in co-administering know analgesics for pain management, to reduce side effects and maximize therapeutic effects by pharmacological synergism, defined as supra-additive effects to biological stimuli. This work aimed to evaluate, using isobolgraphic analysis, synergistic effects of three antinociceptive substances— anandamide (AEA), a cannabinoid CB1 receptor agonist; xylazine (XYL), an adrenergic α_2_-receptor agonist; and DAMGO, an µ-opioid receptor agonist—administered in binary doses in a prostaglandin E2 (PGE_2_)-induced peripheral pain model. Hyperalgesia was induced in Swiss male mice, and subsequently, animals were treated with binary agonist combinations administered to the hind paw. Mechanical nociceptive thresholds were measured using an algesimetric task, and the results obtained were compared with additive predicted effects. For AEA+XYL and AEA+DAMGO combinations, the observed effects were significantly greater than those predicted by Loewe’s additivity principles at all tested effect levels (10%, 30%, and 50% maximum possible effect, MPE). DAMGO+XYL combination showed significant synergistic effects at 10% and 30% MPE but not at 50% MPE. Confirming these findings, combination indexes (CI) for AEA+XYL and AEA+DAMGO were less than 1, indicating synergism, while CI for DAMGO+XYL was near 1, indicating additivity. Notably, single-system antagonism with either AM251, a CB1 antagonist, yohimbine, an α_2C_-receptor antagonist or naloxone, pan-opioid receptor antagonist, could prevent synergy or any analgesia at all for AEA+XYL and AEA+DAMGO. Furthermore, the binary agonist combinations did not produce systemic effects, sedation, or motor impairments. The results suggest synergistic antinociceptive effects for AEA+XYL and AEA+DAMGO, which are dependent on concomitant agonism upon known metabotropic receptors.

## 1. Introduction

Despite the ubiquitous presence of pain among individuals, affecting 30% of the adult population, and the considerable amount of people reaching healthcare services seeking treatment, the pharmaceutical repertoire to effectively manage pain is still limited (Alorfi, 2023; Cohen et al., 2021). Moreover, current analgesics present a multitude of side effects, justifying the need for new approaches (Labianca et al., 2012). Combining existing drugs is considered a promising tool to approach pain, with less cost but good efficacy and safety. Co-administration of existing analgesics is a good method to maximize therapeutic effects with concurrent reduction of side effects (Varrassi et al., 2017). However, there is still limited evidence supporting synergistic effects of combining analgesics for peripheral hyperalgesia.

Pharmacological synergism is a property explored when drugs are combined to achieve a desired supra-additive effect (Grabovsky and Tallarida, 2004). Despite the intuitive comprehension of synergy as pharmacological cooperation, quantitative approaches must rely on clear measurements, which must relate to dose-response relationships for separate and joined drugs, for assessment of supra-additive effects (Geary, 2013; Lederer et al., 2019). Based on Loewe’s Additivity Principle, synergism measurements are feasible in the scope of isobolographic analysis (Loewe, 1928), an analytical and graphical technique to determine dose-response relationships departing from an ideally additive effect for drug combinations (Tang et al., 2015). The method relies on principles of equi-effective doses for combined drugs without any prior mechanistic knowledge and is a golden standard for synergism assessment. Isobolographic analysis helps to determine the synergistic, additive, or antagonistic interactions between drugs, thereby, optimizing combination therapies (Tallarida, 2012, 2011, 2006).

Evidence points out toward the crucial role of peripheral endogenous modulators, such as endocannabinoids, opioids, and catecholamines, in controlling noxious inputs before they are centrally processed (Martínez and Abalo, 2020; Piomelli and Sasso, 2014). Along with their receptors and metabolizing enzymes, these substances determine a tonic control of peripheral nociception, maintained by cannabinoidergic, opioidergic, and adrenergic endogenous systems (Hong and Abbott, 1995; Romero et al., 2013, 2012; Romero and Duarte, 2009). Most existing isobolographic analyses tend to focus on more traditional painkiller combinations or single-class drug interactions in thermal, inflammatory and neuropathic pain models (Delgado et al., 2023; Fischer, 2012; Hurley et al., 2002; Kim et al., 2022; Miranda et al., 2014, 2006), placing a gap in understanding the synergistic benefits of combining agonists in the hyperalgesia pain models. Moreover, the mechanisms behind synergistic interactions of cannabinoidergic, adrenergic, and opioidergic agonists are not completely understood.

The present work is a rational approach based on isobolographic analysis to address the systematic combination effects of cannabinoidergic, adrenergic, and opioidergic agonists and the possible mechanism behind their synergistic interactions. Using a rapid and precise assessment of antinociceptive interactions between three pharmacological agonists – anandamide (AEA), xylazine (XYL), and DAMGO – combined in binary dose regimens, this study offers new insights into potential therapeutic strategies in peripheral pain. Moreover, the exploration of these specific combinations provided a straightforward and statistically robust methodological landscape transposable to any binary analgesic combination using a mouse model of nociceptive pain.

## 2. Methods

### 2.1. Animals

Male Swiss mice, weighing 35–40g, were housed in groups of four/five, with free access to food and fresh water, and kept under controlled temperature (23 ± 1 °C) and 12 h light/dark cycle. Two days prior to all experiments, animals were acclimated to behavioral assay conditions to acquaint themselves with the paw withdraw task or rotarod test. At the end of the experimental procedures, all animals were euthanized with intraperitoneal injection of anesthetic lethal doses (ketamine, 300 mg.kg^-1^ and xylazine, 15 mg.kg^-1^, both Sigma-Aldrich, USA). The study was approved by the Committee of Ethics in Animal Use of the Federal University of Minas Gerais, protocol 69/2018, and followed international guidance for proper animal handling and experiments.

### 2.2. Drugs

The cannabinoid agonist Arachidonoylethanolamide (Anandamide, AEA, Tocris, EUA), adrenergic agonist Xylazine Chloride (XYL, 2% m.V^-1^, Anasedan®) and opioid agonist Ala^2^-MePhe^4^-Gliol^5^-encephalin (DAMGO, Tocris) and their specified doses were diluted to a final injection volume of 20µL either in sterile NaCl 0,9% m.V^-1^ (saline, SLN) or in Tocrisolve®, Tocris (TS). The cannabinoidergic (AM251, Tocris), α_2C_-adrenergic (yohimbine – YOH, Sigma) and opioidergic (naloxone – NLX, Sigma) antagonists were dissolved or diluted in SLN. Prostaglandin E_2_ (PGE_2_) (Sigma, USA) was diluted in ethanol, then in saline obtaining 10% final concentration of alcohol. All solutions were prepared immediately prior to the experiments. Drugs were administered subcutaneously on the plantar surface of the hind paws (volume of 20 μL/paw). The dose of each drug used in this study was previously determined by other works published by our research group (Da Fonseca Pacheco et al., 2008; Romero et al., 2013, 2012; Romero and Duarte, 2009).

### 2.3. Behavioral tests

#### 2.3.1. Nociceptive threshold assessment – paw compression method

The peripheral nociceptive threshold was measured by the mechanical paw pressure test submitted to blunt pressure, as previously described (Kawabata et al., 1992; RANDALL and SELITTO, 1957). Briefly, hyperalgesia was induced by intraplantar (i.pl.) injecting 20 µL of PGE_2_ 0.1 ug.uL^-1^ in the right hind paw at t = 0 min. AEA (12.5, 25 and 50 ng), XYL (25, 50 and 100 µg) and DAMGO (0.25, 0.5, 1 and 2 µg) were injected i.pl. into the right hind paw at t = 175 min. At t = 180 min, animals were subjected to an assessment of the mechanical nociceptive threshold, which involves applying increasing pressure in their treated paw by a flat circular tip positioned facing the plantar surface. Pressure increment was related to the advance of a 10.0 g halter through a ruler, in which each advanced centimeter corresponds to 560 kPa added pressure to the plantar surface (10 g.cm^-1^). When the animal withdrew its paw in response to the increasing pressure, the value in the ruler was taken as an experimental readout and indicated the nociceptive threshold.

#### 2.3.2. Cross antagonism experiment

To assess how cannabinoidergic (AM251), α_2C_-adrenergic (YOH) and opioidergic (NLX) single antagonism affects peripheral antinociception of binary combinations AEA + XYL and AEA + DAMGO, cross-antagonism experiments were performed. For AEA + XYL, PGE_2_ 2 µg was injected i.pl in the left hind paw of animals at t = 0 min. At either t = 145 min (YOH, 20 µg) or t = 165 min (AM251, 80 µg), antagonists were injected i.pl, and at t = 175 min, AEA 16 ng + XYL 16 µg were administered i.pl. Then, at t = 180 min, animals were subjected to the algesimetric task. For AEA + DAMGO, PGE_2_ 2 ug was injected i.pl at t = 0h; at t = 145 min (NLX, 50 µg) or t = 165 min (AM251, 80 µg), the antagonists were injected, and, at t = 175 min, the animals received AEA 14ng + DAMGO 0.1µg i.pl, and finally at t = 180 min the algesimetric task was performed.

#### 2.3.3. Evaluation of systemic effects

Systemic antinociception exclusion was performed, treating both the right and left hind paws i.pl. with the described PGE_2_ dose at t = 0h. At t = 175 min, only the right hind paw was injected with agonists in the highest binary doses used in the experiments. At t = 180 min, both paws were subjected to the withdrawal test, and any statistically significant differences between the mean threshold measured for the left paws and the negative hyperalgesia controls were assumed to be due to systemic antinociception.

#### 2.3.4. Rotarod test

Motor performance and sedation of mice were evaluated using the rotarod test. Animals underwent daily training in the test for two consecutive days before the experiments commenced. The highest binary doses used in the experiments, or their vehicles were injected i.pl. in the right hind paw of animals at t = 0h. After 5 minutes, the animals were subjected to the referred test, and their latency in the rolling bar was compared with negative and positive (XYL 16mg.kg^-1^ intraperitoneal) controls. A trial ended if the mice fell off the drum or the time reached 2 min.

### 2.4. Isobolographic analysis

First, dose-response relationships were interpolated for AEA, DAMGO, and XYL, with ascending doses being administered through i.pl. injection. Response saturation was assumed for total reversion of hyperalgesia, i.e., no difference between the treatment and naïve animals. Responses were expressed as a percentage of Maximum Possible Effect (% MPE), calculated as:

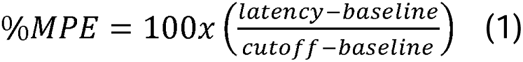

in which the established cutoff was 160 g (to avoid tissue damage in paws) and the baseline corresponded to the mean latency of untreated hyperalgesia controls for each experiment (g). Moreover, those negative controls were assumed to conform to a dose of drug equal to zero. The relationships obtained were fitted to a general four-parameter model to account for the steepness of the curves, as they are better described if Hill’s coefficient is not set to 1. The number of tested doses was kept to a minimum, which allowed good fitting parameters, reducing the number of animals used but preserving pharmacological sense. The fitting equation was:

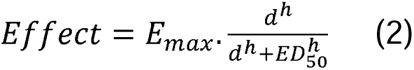

in which *E_max_* corresponds to *% MPE* at saturation (upper plateau), *d* is the given dose of agonist, *h* is the Hill slope measuring the steepness of the dose-response relationship, and *D_50%_*is the dose in which *effect* = 50% *E_max_*. Curve fitting was performed using the software GraphPad Prism 8.0. The goodness of fit was assumed for at least R^2^ ≥ 0.85.

After obtaining the dose-response relationships, additive isoboles were constructed for binary combinations of agonists using the set of parameters derived from the interpolated model. No linearity for isoboles was assumed since all agonists displayed distinct potencies and efficacies (Grabovsky and Tallarida, 2004). Briefly, for a binary combination of agonists A and B, additive isoboles for 10%, 30%, and 50% MPE were given as a function *b = b(a)*, defined in the upper-right quadrant of a Cartesian plane (dose plane), in which *a* and *b* represents an ordinate pair *(a,b)* of doses of agonists A and B, respectively:

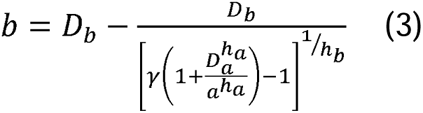

in which *D_b_* and *D_a_* are the respective doses for E_10_, E_30,_ and E_50_ for B and A, γ is the ratio between *E_max_*for B and *E_max_* for A, and *h_a_* and *h_b_*are Hill coefficients for each agonist A and B, respectively. Theoretical additive points in each isobole were determined for a fixed dose ratio (doses in µg) of agonists, and for reducing the animal number, only one fixed dose ratio was chosen for each binary combination: 100:1 for AEA/XYL, 50:1 for AEA/DAMGO and 30:1 for DAMGO/XYL. Intersection points were tested in the paw withdraw test to determine the experimental effect (E_a,b,Experimental_). All the points tested experimentally are described in Table 1.

**Table 1.** Doses used in the binary agonist combinations for 10, 30, and 50% MPE.

| Binary Combination | %MPE | AEA/ng | XYL/μg | DAMGO/μg |
| --- | --- | --- | --- | --- |
| AEA + XYL | E<sub>10</sub> | 7.5 | 7.5 | - |
|  | E<sub>30</sub> | 12 | 12 | - |
|  | E<sub>50</sub> | 16 | 16 | - |
| AEA + DAMGO | E<sub>10</sub> | 6 | - | 0.12 |
|  | E<sub>30</sub> | 9 | - | 0.18 |
|  | E<sub>50</sub> | 14 | - | 0.28 |
| DAMGO + XYL | E<sub>10</sub> | - | 6 | 0.20 |
|  | E<sub>30</sub> | - | 12 | 0.40 |
|  | E<sub>50</sub> | - | 18 | 0.60 |

Then, the combination index (CI), defined as a Cartesian metric derived from the binary doses was applied in the isobolographic analysis to compare the theoretical binary dose *(a_theoretical_,b_theoretical_)* for a given effect level with the experimental dose *(a_exp_,b_exp_)* that give such effect) (Chou, 2010). CI is calculated as the ratio between the distances of *(a_exp_,b_exp_)* and *(a_theoretical_,b_theoretical_)* from the origin *(0,0)*, respectively. Deviating values from the unity indicate synergy (CI < 1) or sub-additivity (CI >1). Equation 4 outlines the calculation of CI:

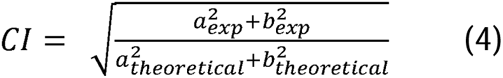

The predicted additive curves for combinations of agonists A and B were established using the following model, in which *b_eq,a_* corresponds to the dose *b* that is equi-effective to a given dose *a*.

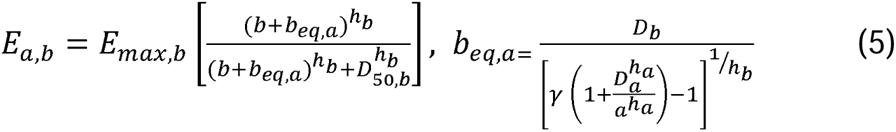

Finally, variances were determined to compare theoretical and experimental effects observed for each binary combination of agonists using a Student’s t-test (Tallarida, 2011). The variance of the effect *V(E_a,b_)* for each theoretical additive point was calculated as:

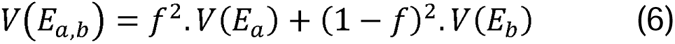

in which *f* is the potency ratio for drugs A and B and *V(E_a_)* and *V(E_b_)* are the variances of the effects of drugs alone.

### 2.5. Statistical analysis

Values were expressed as mean ± SEM (g) or % MPE, and measurements were made in triplicate by two blinded experimenters not involved in data analysis. All data were subjected to parametric analysis (Student’s t-test or one-way ANOVA followed by Bonferroni’s post-hoc test) for pairwise comparison. Significant differences were indicated by p < 0.05. Statistical analyses were performed with Prism 8.0 (Graphpad Software Inc.) and R (R Core Team, 2014).

## 3. Results

### 3.1. Peripheral antinociceptive effects of AEA, XYL, and DAMGO in animals with PGE_2_-induced hyperalgesia

First, the peripheral antinociceptive effects of different classes of agonists were investigated in a dose-dependent manner in the PGE_2_-induced hyperalgesia animal model. AEA, a CB1 agonist, displayed peripheral antinociception in a dose-dependent manner, ranging from 12.5 to 50 ng (p<0.05; One-way ANOVA followed by Bonferroni posthoc test) (Fig. 1A). Similarly, XYL, which activates α_2C_-adrenergic receptors, showed effective antinociception actions at 25, 50 and 100 µg doses (p<0.05; One-way ANOVA followed by Bonferroni posthoc test) (Fig. 1B). Furthermore, the µ-opioid receptor agonist DAMGO displayed dose-dependent peripheral antinociception at 0.25, 0.5, 1, and 2 µg (p<0.05; One-way ANOVA followed by Bonferroni posthoc test) (Fig. 1C). These findings confirm the antinociceptive effects of cannabinoidergic, adrenergic and opioidergic agonist in the pain algesimetric task.

**Figure 1.**
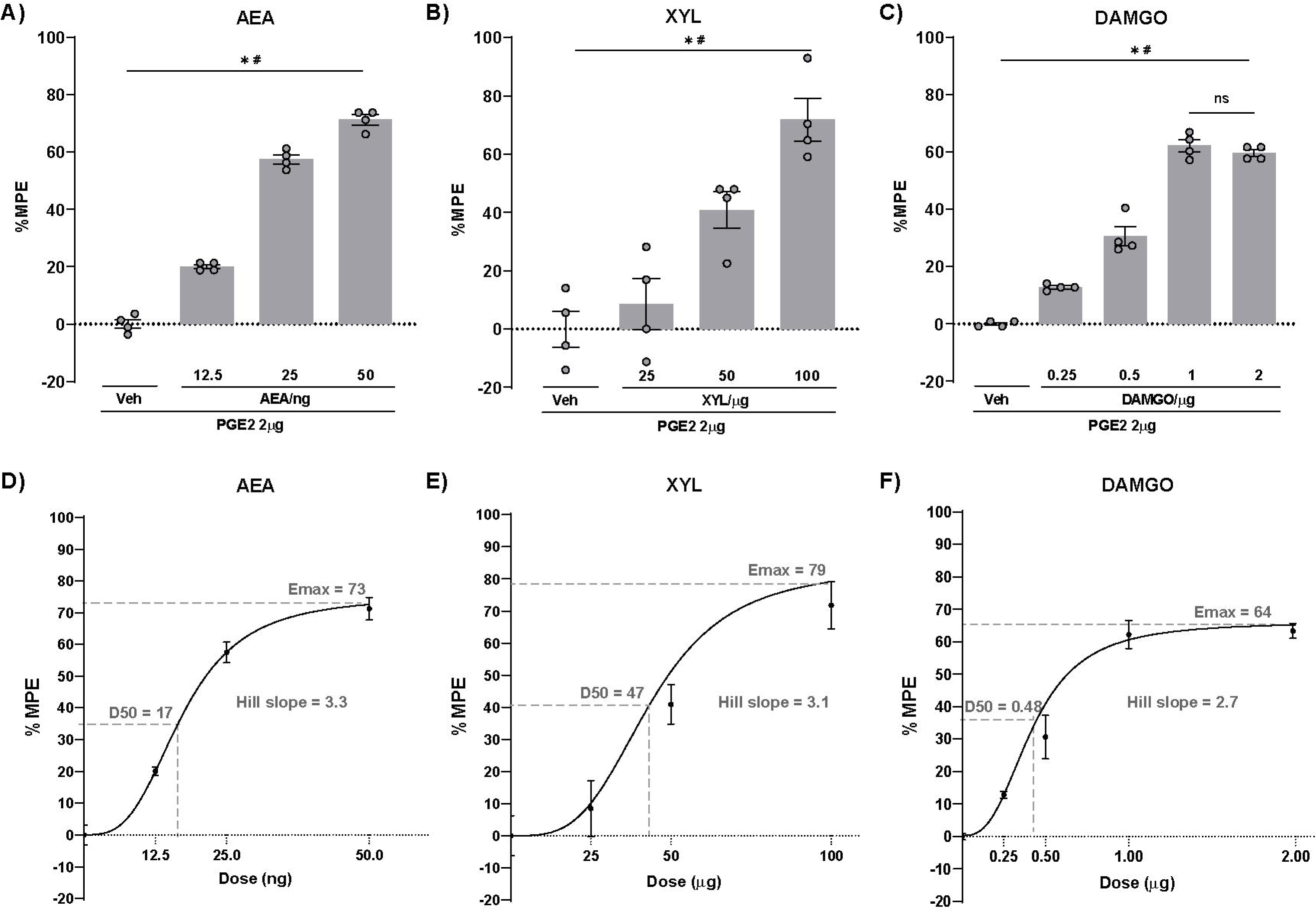
Dose-response curves of peripheral antinociceptive effects in PGE_2_-induced hyperalgesia model. **(A)** % of the maximum possible effect (MPE) of AEA, a CB1 receptor agonist, showing dose-dependent response in the range of 12.5 to 50 ng. All tested doses exhibited significant antinociceptive effects. **(B)** % MPE of XYL, an α_2C_-adrenergic receptor agonist, with effective antinociception at 25, 50, and 100 µg doses. Significant effects were observed at each dose. **(C)** % MPE of DAMGO, a µ-opioid receptor agonist, in a dose-dependent manner at 0.25, 0.5, 1, and 2 µg. Significant antinociceptive responses were observed at each dose compared to vehicle and among them. However, the % MPE did not differ between DAMGO 1 and 2 µg. *p<0.05 indicates significant differences between tested doses and vehicle (tocrisolve or saline) and ^#^p<0.05 indicates significant differences among tested doses in a dose-dependent manner; one-way ANOVA followed by Bonferroni posthoc test. **(D)** Interpolated sigmoidal dose-response curve for AEA, **(E)** XYL, and **(F)** DAMGO. Fitting parameters (E_max_, D_50_, and Hill slope) are described in each graph. Veh = Vehicle; n.s. = non-significant difference.

After interpolating the sigmoidal model for dose-response of AEA (Fig. 1D), XYL (Fig. 1E) and DAMGO (Fig. 1F), the pharmacodynamic antinociceptive profiles observed for all tested agonists revealed a steep (h_AEA_ = 3.3; h_XYL_ = 3.1; h_DAMGO_ = 2.7) and saturable response (E_max.AEA_ = 73% MPE; E_max.XYL_ = 73% MPE; E_max,DAMGO_ = 64% MPE), reinforcing the adequacy of the fitting model. Notably, it was needed to test an extra ascending dose of DAMGO (2 µg) for fitting optimization, as it forced the achievement of an upper plateau in the interpolated curve. Overall, these results support the robustness of the sigmoidal model in acquiring antinociceptive pharmacodynamic profiles of the tested agonists.

### 3.2. Binary combinations of AEA, XYL, and DAMGO produced additive to synergistic peripheral antinociception

All data derived from the fitting model was used to construct theoretical dose-response relationships for a given binary agonist combination (AEA + XYL; AEA + DAMGO; DAMGO + XYL), assuming additivity as a model premise. The theoretical additive effects for all agonist combinations were determined via equation 5, and the binary doses for 10, 30, and 50% MPE were calculated. Three isoboles for each binary combination were constructed, and each point in the isoboles has the given theoretical additive effect represented by the curve. AEA + XYL isoboles were fitted by linear regression (Fig. 2A), and AEA + DAMGO (Fig. 2B) and DAMGO + XYL (Fig. 2C) isoboles were fitted by 2^nd^-order polynomial regression. The goodness of fit was assumed for at least R^2^ > 0,99. Such a repertoire of binary doses was tested experimentally to compare observed antinociceptive responses with the theoretical MPEs chosen. Figures 2A-C depict the theoretical additive effects (circles) for AEA + XYL, AEA + DAMGO, and DAMGO + XYL, along with the experimental data (squares) obtained from the same binary combinations of agonists. For AEA + XYL (Fig. 2A) and AEA + DAMGO (Fig. 2B), all experimental points lie below the isoboles, suggesting synergistic doses. In contrast, theoretical points overlap with experimental points for DAMGO + XYL (Fig. 2C), suggesting additive effects.

**Figure 2.**
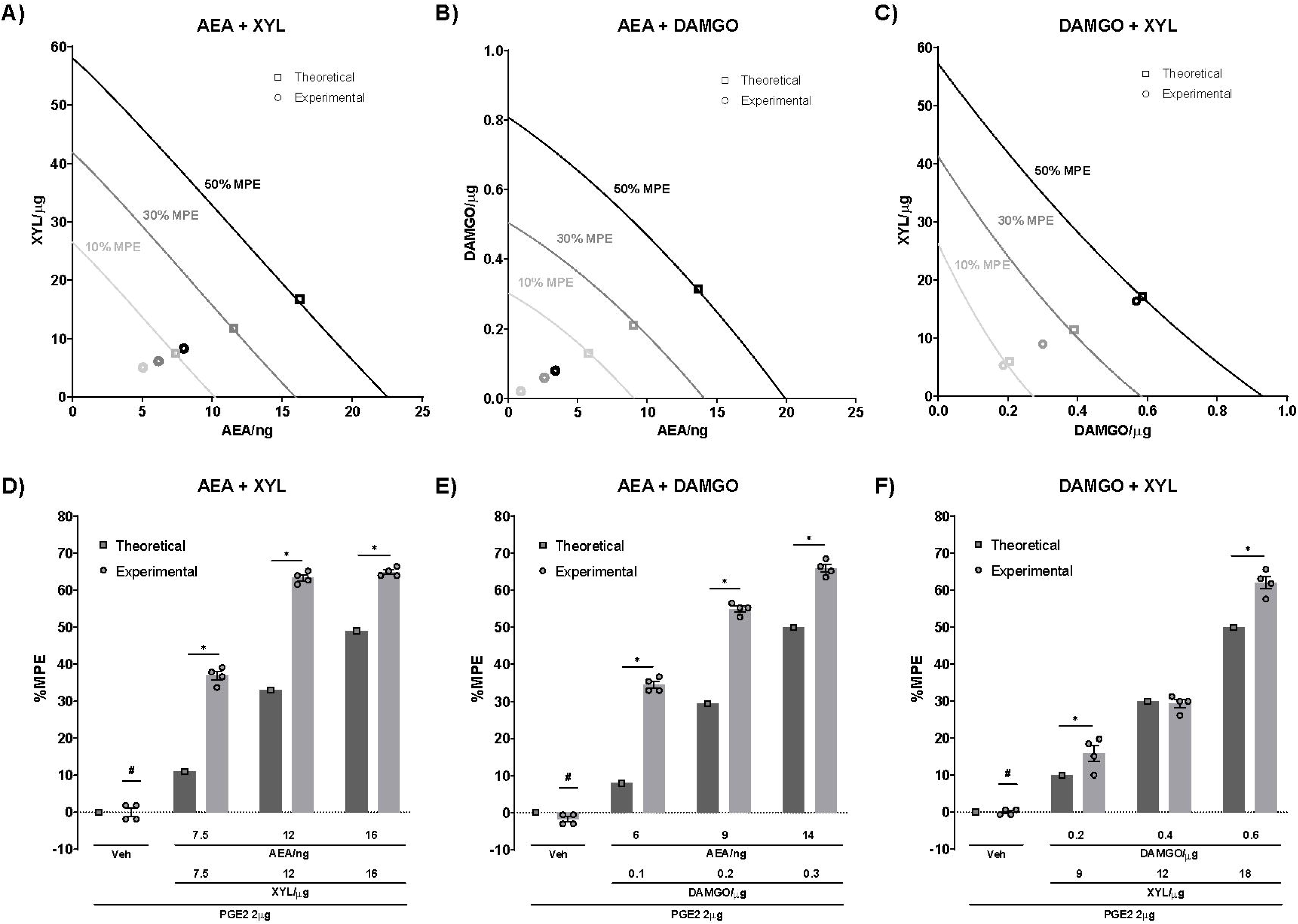
Theoretical and experimental dose-response relationships for binary agonist combinations. Isoboles for the combination of **(A)** AEA + XYL, **(B)** AEA + DAMGO, and **(C)** DAMGO + XYL at 10%, 30%, and 50% maximum possible effect (MPE), fitted by linear regression or 2nd-order polynomial regression. The circles represent the theoretical additive effects, and the squares denote the experimental data. Experimental points below the isoboles indicate a synergistic interaction (AEA + XYL and AEA + DAMGO). Theoretical points overlapping with experimental points imply additive effects (DAMGO + XYL). Comparison of theoretical and experimental data for the **(D)** AEA + XYL, **(E)** AEA + DAMGO, and **(F)** DAMGO + XYL at 10%, 30%, and 50% MPE. Experimental % MPEs were significantly higher than all theoretical predictions, except for 30% MPE for DAMGO + XYL. *p<0.05 indicates significant differences between the theoretical and experimental data; Student’s t-test. #p<0.05 denotes significant differences between vehicle and all experimental tested doses; one-way ANOVA followed by Bonferroni posthoc test. Veh = vehicle (tocrisolve and saline).

To confirm that pharmacological profiles are significantly distinct between theoretical predictions and experimental data, theoretical variances were determined for each binary agonist combination (Tallarida, 2011). A synergistic effect was observed for AEA + XYL (Fig. 2D) and AEA + DAMGO (Fig. 2E), considering that all % MPEs were higher in the experimental approach compared to the theoretical additivity prediction (p<0.05; Student’s t-test). Similar synergistic profiles for DAMGO + XYL were found when 10 and 50 % MPE were compared between theoretical and experimental data (p<0.05; Student’s t-test) (Fig. 2F). However, the 30% of experimental MPE for DAMGO + XYL was not significantly different from the theoretical prediction, indicating an additive antinociception. Overall, these findings demonstrate the varying interaction profiles, i.e., synergistic or additive, among the different binary combinations of cannabinoidergic, adrenergic, and opioidergic agonists in the PGE_2_-induced hyperalgesia model.

Then, combination indexes were calculated from the experimental data obtained to confirm antinociception observed in the binary agonist combinations. Experimental effective doses for a given effect level were derived for each tested point and corresponded to the actual paired dose needed to achieve that given effect level. For instance, considering a dose of AEA + XYL that theoretically gives 30% of the effect level, if synergy occurs, the required dose will be less than the theoretically derived. In this case, the dose will be at the E_9_ (9%) theoretical effect level. Following this rationale, all (a_exp,bexp_) points were calculated for all chosen MPE ratios (Table 2). It can be observed that CI values agree with the effect comparison, being closer to or equal to 1 only for DAMGO + XYL, hence indicating additivity for these binary combinations. For all AEA + XYL and AEA + DAMGO doses tested, the combination acted synergistically, as indicated by values of CI lesser than 1.

**Table 2.** CI of binary agonist combinations for % MPE ratios.

| Binary combination | % MPE ratio | CI |
| --- | --- | --- |
| AEA + XYL | $E_4/E_{10}$ | 0.69 |
| | $E_9/E_{30}$ | 0.52 |
| | $E_{14}/E_{50}$ | 0.48 |
| AEA + DAMGO | $E_1/E_{10}$ | 0.15 |
| | $E_{13}/E_{30}$ | 0.30 |
| | $E_{23}/E_{50}$ | 0.25 |
| DAMGO + XYL | $E_7/E_{10}$ | 1.00 |
| | $E_{25}/E_{30}$ | 0.80 |
| | $E_{52}/E_{50}$ | 1.00 |
| CI = combinatory index; MPE = maximum possible effect. |  |  |

### 3.3. Cross-antagonism mediates the synergistic antinociceptive effects of binary agonist combinations

To further investigate mechanistic features involved in the synergistic analgesia elicited by AEA + XYL and AEA + DAMGO, pharmacological antagonism experiments were performed, in which a single antagonist was used against a binary dose of agonists. The CB1 antagonist AM251 (80 µg), when administered i.pl prior to both AEA 16 ng + XYL 16 µg (Fig. 3A) and AEA 14 ng + DAMGO 0.3 µg (Fig. 3B) treatments fully blocked any analgesia putatively associated to the agonists (p<0.05; one-way ANOVA followed by Bonferroni post-hoc test), even though the doses used were supposed to elicit a theoretical 50 % MPE. The α_2C_-adrenergic antagonist YOH (20 µg), when administered i.pl before AEA 16 ng + XYL 16 µg, also abolished any analgesia attributable to the agonists (Fig. 3A), as well as the pan-opioid antagonist naloxone (50 µg), which also impairs any analgesic effect of AEA 14 ng + DAMGO 0.3 µg (Fig. 3B), when administered i.pl before the agonists (p<0.05; one-way ANOVA followed by Bonferroni post-hoc test). These results reinforce the view of synergistic peripheral analgesia mediated by cannabinoidergic/adrenergic and cannabinioidergic/opioidergic systems, as single-system pharmacological blockage abolishes all agonist-mediated antinociception, in spite of their distinct classes.

**Figure 3.**
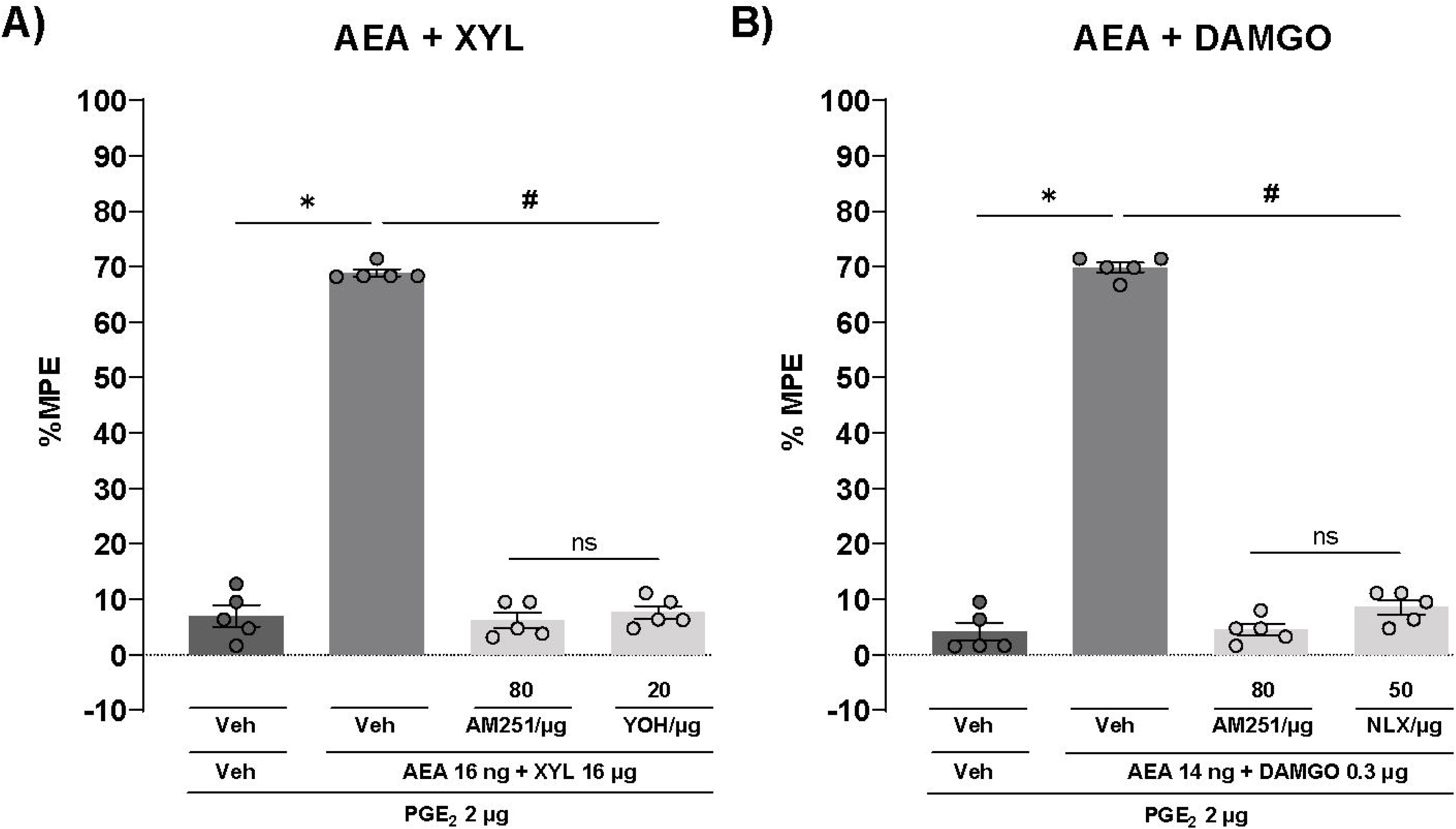
Effects of cannabinoidergic, adrenergic and opioidergic antagonism on peripheral antinociception induced by binary agonist combinations in a PGE_2_-induced hyperalgesia model. **(A)** The peripheral antinociceptive effect of AEA (16 ng) combined with XYL (16 µg) was reversed by AM251 (a CB1 receptor antagonist) and Yohimbine (YOH, an α2c-adrenergic antagonist), as shown by the % maximum possible effect (MPE). **(B)** Similarly, the % MPE of AEA 14 ng + DAMGO 0.3 µg in the hyperalgesia model was reduced after administration of AM251 or naloxone (NLX, µ-opioid antagonist receptor). *p<0.05 indicates significantly difference between tested binary agonist combinations and vehicle (Trocrisolve and saline) and #p<0.05 denotes differences between tested doses and antagonists; one-way ANOVA followed by Bonferroni posthoc test. Veh = vehicle; n.s. = non-significant difference.

### 3.4. The binary agonist combinations did not produce any systemic effects, sedation, or motor impairments

To preclude any possible systemic analgesia due to binary agonist combinations, PGE_2_ (2 μg/paw) was administered to both hind legs of the animals, and tested doses were injected i.pl. into the right paw while saline was administrated into the left paw. The antinociceptive effect of AEA 16 ng + XYL 16 µg, AEA 14 ng + DAMGO 0.3 µg, and XYL 17 µg + DAMGO 0.6 µg were confined to the treated paw (p<0.05; one-way ANOVA followed by Bonferroni post-hoc test), indicating local action without central pathway involvement (Fig. 4A). Additionally, in the rotarod test, no significant differences were observed in the treated groups AEA 16 ng + XYL 16 µg, AEA 14 ng + DAMGO 0.3 µg, and XYL 17 µg + DAMGO 0.6 µg when compared with control groups (Fig. 4B), indicating that binary agonist combinations at maximum doses did not produce sedation or motor impairments. The positive control group, XYL 500 µg, decreased rotarod latency to fall off, thereby confirming the expected pharmacological effect of motor impairment (p<0.05; one-way ANOVA followed by Bonferroni post-hoc test).

**Figure 4.**
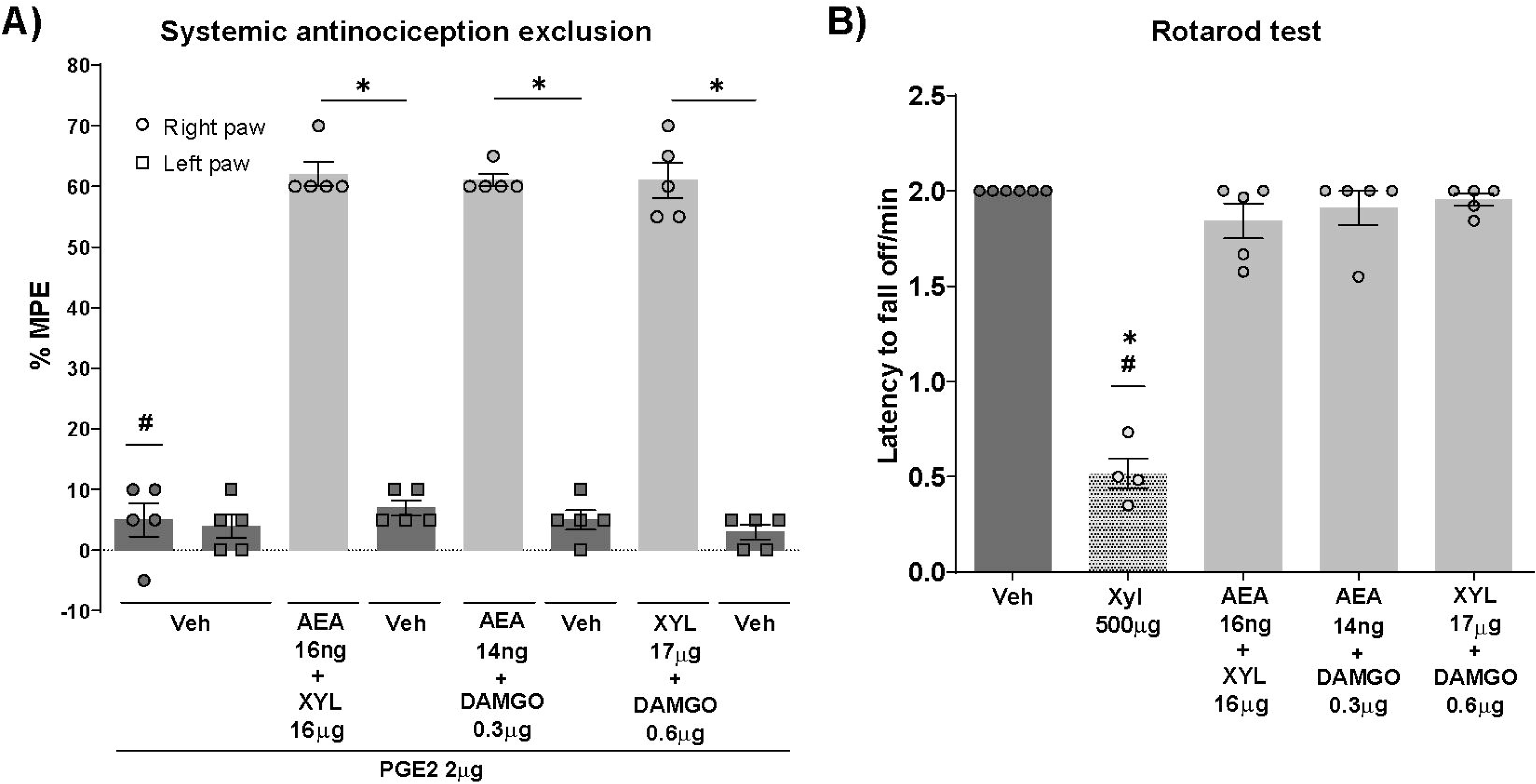
Local antinociceptive effects and motor coordination analyses for binary agonist combinations. **(A)** Antinociceptive effects were exclusively observed in the right hind paw, treated with the maximum doses of binary agonist combinations: AEA 16 ng + XYL 16 µg, AEA 14 ng + DAMGO 0.3 µg and XYL 17 µg + DAMGO 0.6 µg compared to the left hind paw injected with vehicle (Tocrisolve and saline), indicating localized action. *p<0.05 indicates significant differences between the tested doses injected in the right hind paw and vehicles injected in the left hind paw and #p<0.05 vehicle vs. tested doses injected in the right hind paw; one-way ANOVA followed by Bonferroni posthoc test. **(B)** No significant differences were found between the treated groups (AEA 16 ng + XYL 16 µg, AEA 14 ng + DAMGO 0.3 µg, and XYL 17 µg + DAMGO 0.6 µg) and control groups in the rotarod test, indicating no motor impairment. The positive control group (XYL 500 µg) showed decreased latency to fall when compared to vehicle and tested doses. *p<0.05 XYL 500 µg vs. vehicle; #p<0.05 XYL 500 µg vs. tested doses; one-way ANOVA followed by Bonferroni posthoc test. MPE = maximum possible effect. Veh = vehicle.

## 4. Discussion

Considering the persistent prevalence of pain in the adult population, the existing pharmacological treatments’ limited efficacy, and the substantial side effects of painkillers, the combination of current analgesics offers a less expensive and possibly more effective option (Huang et al., 2019; Varrassi et al., 2017). This study confirmed the individual efficacy of AEA, XYL, and DAMGO in producing dose-dependent antinociceptive effects in a PGE_2_-induced hyperalgesia model. Using isobolographic analysis, AEA + XYL and AEA + DAMGO displayed synergistic interactions, while XYL + DAMGO showed additive effects in the proposed model. Notably, high doses of binary agonist combinations did not produce systemic antinociception and motor impairment, supporting the safety of these combinations. Furthermore, it was observed that a single class of antagonists can reverse the synergistic antinociceptive effects of binary agonist combinations, shedding light on the underlying mechanisms of their interactions.

AEA, XYL, and DAMGO are described as ligands of representative G-protein coupled receptors (GPCRs) of the cannabinoidergic, adrenergic, and opioidergic system, respectively, eliciting a classical G_i/o_-mediated response (Di Marzo et al., 2000; Törneke et al., 2003; Zöllner et al., 2003). Activation of G_i_ receptors leads to desensitization of primary sensory neurons and results in downstream cell signaling, including lower levels of intracellular cAMP, increased K^+^ conductance (hyperpolarization), and reduced Ca^2+^ conductance (diminished Ca^2+^-dependent exocytosis of synaptic vesicles) (Stein et al., 2009). Indeed, in this study, AEA, XYL, and DAMGO agonists alone were capable of eliciting antinociception in a dose-dependent manner in the PGE_2_-induced hyperalgesia model, confirming the previous findings from our group (Romero et al., 2020, 2013, 2012; Romero and Duarte, 2009). AEA, an endogenous cannabinoid that acts as a partial CB1 receptor agonist, has therapeutic potential for managing a variety of types of pain, including thermal and chemical hyperalgesia and neuropathic and cancer pain (Desroches et al., 2014; Khasabova et al., 2013; Schreiber et al., 2012). XYL, an α_2_-adrenergic agonist commonly used as a sedative and muscle relaxant in veterinary medicine, can modulate pain perception by reducing norepinephrine release (Kitano et al., 2019). DAMGO binds with high affinity to the µ-opioid receptor and is a synthetic analog of endogenous opioids with a strong morphine-like antinociceptive effect (Al-Khrasani et al., 2007; Mecklenburg et al., 2017). Furthermore, the AEA, XYL, and DAMGO antinociceptive profiles obtained in this study fit with good concordance with a four-parameter model, in which the effects were clearly saturable with steep correlation, indicating robust antinociceptive effects in the PGE_2_-induced hyperalgesia model.

Assessments of synergy for binary mixtures of drugs are based on Loewe’s Additivity Principle and rely strictly on dose-response relationships for individual substances. Such dose-effect relations are founded on the Law of Mass Action and the concept of pharmacological receptors, dictating in conjunction with the adequate fitting model to be used (Tallarida, 2012). In this study, the combinations of doses for synergism assessment were established for 10, 30, and 50% MPE, as determined by the intersection of fixed-dose ratio curves with the isoboles. These lines on the dose plane behave, in a pharmacological sense, as drugs themselves (Gessner, 1995), with theoretical parameters determinable from an additive perspective. It is noted that the effects of AEA + XYL and AEA + DAMGO were significantly greater than the theoretical assumption of additive effects, in contrast with the pair XYL + DAMGO at the 30% MPE. In line with the isobolographic analysis, the combination indexes (CI) for AEA + XYL and AEA + DAMGO were below 1, indicating synergism. However, while the effect analysis suggested a synergism for 10% and 50% MPE for XYL + DAMGO, CI values for all binary combinations were near 1. Such divergent results for XYL + DAMGO may be explained by low standard errors resulting from precise measurements in the algesimetric task. Hence, it is crucial to consider both isobolographic and CI analyses for accurately assessing and defining synergy between doses. Overall, these results indicate that AEA + XYL and AEA + DAMGO combinations act synergistically as antinociceptive agents in the PGE_2_-induced hyperalgesia model.

Using isobolographic analysis in thermal analgesia assays, synergistic interaction between CP55,940, a synthetic cannabinoid agonist, and morphine was observed in both tail flick and hot plate tests (Tham et al., 2005). In a mouse neuropathic pain model, the pan-cannabinoid receptor agonist WIN55,212-2 combined with morphine synergistically reduced allodynia (Kazantzis et al., 2016). Furthermore, the co-administration of morphine with agonists of cannabinoid receptors induced synergistic thermal antinociception in diabetic animal models and mechanical analgesia in arthritic pain models, as well as reduced inflammatory pain (Auh et al., 2016; Cox et al., 2007; Williams et al., 2008; Yuill et al., 2017). These findings support that the combination of AEA and DAMGO in doses predicted by isobolographic analysis elicits a synergic effect in the peripheral hyperalgesia model. Concerning co-administration of cannabinoid and adrenergic agonists, CP55,940 and dexmedetomidine exhibited antinociceptive synergy in the hot plate assay (Tham et al., 2005), as observed for AEA + XYL in this study. However, in the tail-flick assay, CP55,940 and dexmedetomidine showed additive outcomes, underscoring that the effects of drug combinations can vary depending on the specific type of pain assessment. Consistent with the additive effects of XYL + DAMGO in the PGE_2_-induced hyperalgesia model, the combination of dexmedetomidine and morphine resulted in additive effects in the hot plate and tail-flick assays (Tham et al., 2005). Notably, the administration of adrenergic and opioid receptor agonists in binary combinations via intrathecal results in antinociceptive synergy in a variety of pain models (Fairbanks et al., 2000a, 2000b; Przesmycki et al., 1997). Therefore, it is important to point out divergent synergistic central and peripheral antinociception mediated by the combination of adrenergic and opioid agonists.

Even though it is not possible to assess any mechanistic explanations for the observed synergistic antinociception between AEA + XYL and AEA + DAMGO based on isobolographic analysis, the agonist properties of these substances upon classical inhibitory GPCRs may offer some highlights. Considering the co-expression of cannabinoid, adrenergic, and opioid metabotropic receptors on the same neuron, their shared intracellular signaling pathways may co-potentiate activation by ligands, possibly by enhancing the pool of intracellular partners needed to engage to active receptors for signal transduction, as G-proteins, towards the other receptor, and vice-versa (Djellas et al., 2000). Another possibility pertaining to the co-expression of those GPCRs on a given neuron is the occurrence of receptor heterodimers, which are supramolecular GPCR assemblies that possess unique signaling features, especially when heterodimeric partners are regarded as true allosteric modulators upon each other (Haack and Mccarty, 2011). Indeed, this is extensively shown to µ-opioid receptors and several other partners, including CB1 and α_2_-adrenergic receptors, and the signaling consequences of MOR heterodimerization are evaluated up to behavior (Zhang et al., 2020).

Antinociceptive effects of AEA, XYL and DAMGO are readily blocked pharmacologically using classic GPCR antagonists, a paradigm extensively explored in experimental therapeutics targeting pain management that reveals a strong receptor-dependent antinociceptive effect of such substances (Romero et al., 2020, 2013, 2012; Romero and Duarte, 2009). To capture the dependency of observed synergistic analgesic effects of the binary doses AEA + XYL and AEA + DAMGO at the receptor level, it was sought to antagonize a single endogenous system while activating both tested. The CB1 antagonist AM251, known for blocking peripheral analgesia mediated by AEA (Romero et al., 2013), showed to suppress synergy between AEA + XYL and AEA + DAMGO, but also to block any analgesia from XYL or DAMGO, respectively. It is interesting that the same trend was observed for YOH in the AEA + XYL co-administration regimen and for NLX in the binary combination AEA + DAMGO. These results are even more relevant if considered the disparate molecular structures of AM251, YOH and NLX, as well as the distinct orthosteric sites for the CB1, α2C-adrenergic and µ-opioid receptors (Huang et al., 2020; Serohijos et al., 2011; Wu et al., 2021). Such divergence in structure and affinity for distinct receptors rules out cross-blockage, by the antagonists used, of targeted GPCRs, revealing that receptor downstream signaling is relevant for the synergistic analgesia observed. Moreover, the total lack of analgesia by the non-blocked agonist partner in the binary doses brings a role to the receptors themselves are players in this finely tuned crosstalk. The pharmacological antagonism of one system cannot be compensated by other agonism upon a second system, if there is any agonism at all, revealing overlapping and balanced interplay between pathways.

The peripheral action of the binary doses used was confirmed by systemic exclusion experiments in which both systemic analgesia and sedation were discarded. The lack of systemic analgesia is a plus for such a kind of combinatorial experiment, confirming a peripheral mechanism to circumvent pain. It is noteworthy to point out that sedation is also a side effect associated with analgesia (PHILLIPS et al., 2017), and the lack of sedation and motor impairments, as assessed by the rotarod task, indicates that these combinations do not appear to scale side effects. Aligned with these findings, when the synthetic cannabinoid agonist WIN5512 was combined with morphine in a neuropathic pain model, no cataleptic phenotype was observed in the bar test or sedation-like phenotype in the open field test (Kazantzis et al., 2016). These results emphasize the potential of such combinatorial approaches to provide effective pain relief without undesirable side effects.

## 5. Conclusions

Here it is provided evidence for synergistic peripheral analgesia mediated by endogenous cannabinoidergic/adrenergic and cannabinoidergic/opioidergic systems, when these are concomitantly activated by known agonists administered in binary combinations, namely AEA+XYL and AEA+DAMGO, respectively. The detailed isobolographic analysis conducted reveals strong analgesic synergy for the cited binary combinations, a feature that reinforces the adequacy of clinical efforts to combine analgesics of distinct classes to reduce doses and increment therapeutic outcomes. Notably, single-system antagonism prior to the binary doses administration was shown to prevent not only synergy, but any analgesia at all, pointing to a complex interplay between these endogenous systems in modulating peripheral pain. Such a complexity, provenly emergent by being synergistic and dependent on input (concomitant agonism) offers an interesting avenue for exploring combinatorial pharmacology of analgesics at both pre-clinical and clinical levels, a trend to be better explored.

## DECLARATION OF INTEREST

The authors declare that they have no conflict of interest concerning the subject of this study.

## CRediT AUTHORSHIP CONTRIBUTION STATEMENT

**C.F.B.L.:** writing (original draft, review and editing), methodology, investigation, formal analysis, data curation, conceptualization; **T.S.S.:** writing (original draft, review and editing), methodology, investigation. **F.C.F:** writing (review and editing), methodology, investigation. **B.F.G.Q.:** methodology, investigation. **I.D.G.D.:** writing (review and editing), validation, project administration, funding acquisition. **T.R.L.R.:** writing (review and editing), validation, supervision, project administration, funding acquisition, data curation, conceptualization.

## FUNDING

This work received financial support from the National Council for Scientific and Technological Development (CNPq) – Brazil (MCTI/CNPQ/Universal 14/2014, number 448283/2014-0). TRLR (Productivity Fellowship, Level 2, number 301496/2018-8) and IDGD (Productivity Fellowship, Level 1D, number 301496/2018-8) thank the financial support from CNPq. CFBL, TSS, FCF, and BFGQ acknowledge fellowships from Coordination for the Improvement of Higher Education Personnel (CAPES), Brazil.

## DATA AVAILABILITY STATEMENT

The data underlying this article are available in the article and its online supplementary material.

## Notes

### Competing Interest Statement

The authors have declared no competing interest.

